# Performance evaluation of structural variation detection using DNBSEQ whole-genome sequencing

**DOI:** 10.1101/2024.11.28.625961

**Authors:** Junhua Rao, Huijuan Luo, Dan An, Xinming Liang, Lihua Peng, Fang Chen

**Affiliations:** MGI Tech, Shenzhen 518083, China; BGI, Shenzhen 518083, China

## Abstract

DNBSEQ platforms have been widely used for variation detection, including single-nucleotide variants and short insertions and deletions, comparable to Illumina. However, the performance of structural variations (SVs) detection using DNBSEQ platforms remains unclear. We assessed SV detection using 40 tools on eight DNBSEQ and two Illumina NA12878 whole-genome sequencing datasets. We confirmed that the performance of SV detection on DNBSEQ and Illumina platforms is consistent when using the same tools, with high consistency in metrics of number, size, precision, and sensitivity. Furthermore, we integrated representative SV sets for both DNBSEQ (4,785 SVs) and Illumina (6,797 SVs) platforms. We found high consistency between the DNBSEQ and Illumina SV sets in terms of genomic characteristics, including repetitive regions, GC distribution, difficult-to-sequence regions, and gene features, indicating the value of both platforms in understanding repetitive regions and the genomic context of SVs. Our study provides a benchmark resource for further studies of SVs using DNBSEQ platforms.

## Introduction

Structural variation (SV) is a general term for different types of genomic mutations with size large than 50bp, including deletion (DEL), insertion (INS), duplication (DUP), inversion (INV) and translocation (TRA)^1^. DELs and DUPs are also classified as copy-number variations (CNVs)^1^. SVs differ from small variants, such as single-nucleotide variants (SNVs) and short insertions and deletions (INDELs), in size and formation mechanisms^2^. SVs significantly contribute to the diversity found within human populations and have a notable impact on human health and disease^3, 4, 5, 6, 7^. In recent years, the importance and mechanism of SVs in human populations and diseases has been further explored and confirmed through large-cohort studies^8, 9^. The 1000 Genomes Project (1KGP) analyzed SVs of 2,504 individuals and estimated that SVs were ∼50-fold enriched for expression quantitative trait loci (eQTLs) compared with SNVs^6^. The Human Genome Structural Variation Consortium (HGSVC) identified 107,590 SVs with 278 SV hotspots in human genome and explored the contribution of SVs in population adaptive selection of humans^10^. Collins et al. constructed 163 genome-wide dosage sensitive segments of rare SVs across 54 disorders for human disease searches^11^.

Currently, several large-cohort studies investigating human disease or human population described above are based on short-read whole-genome sequencing (WGS) technology^6, 8, 9, 12^. This is attributed to the development and utilization of massively parallel sequencing (MPS) platforms that generate short-read data, along with the advancements in analytical tools for detecting SVs. Dozens of tools that can use short-read (50-150 bp) WGS data sequenced on MPS platforms to detect SVs on a genome-wide scale^13^. Each SV tool is based on one of the following five algorithms: (1) read depth (RD), (2) read pair (RP), (3) split read (SR), (4) de novo assembly (AS) and (5) combination of approaches (CA). Therefore, the types of SVs detected by each tool may different. For instance, tools based on RD algorithm can only detected DELs and DUPs, such as CNVnator^14^. Some tools are specifically designed to detect only specific types of SVs, such as BASIL-ANISE^15^ for INSs and Sprites^16^ for DELs. Meanwhile, the sensitivities of SVs were reported to fluctuate in the range of 10%-70% depending on the size and type of SVs, while the false-positive rates were up to 89%^17^. Despite these limitations, SV detection on short-read data sequenced on MPS platforms is still a good approach for SV research due to variables such as cost, time, resolution and project scope^6, 9^. In short, the SV tools described above were mainly designed to detect SVs based on datasets from Illumina platforms, and the performance of SVs detected by these SV tools has been reported in many articles^13, 18,19^.

As is known to all, Illumina platforms, such as HiSeq 2500 and NovaSeq6000, are the main MPS platforms widely used in research of SVs^12^. For example, leveraging ∼30X WGS data generated by the NovaSeq6000 system, researchers have broadened the spectrum of genomic variants, including the SV catalog, for the 1KGP^6, 9^. Since 2015, the DNBSEQ sequencing platforms, based on the technologies of DNA nanoballs (DNBs) and combinatorial Probe-Anchor Synthesis (cPAS), has been widely utilized in genomic researches for its high sequencing accuracy, low duplication rates, and reduced index hopping^20^. To date, the DNBSEQ platforms have been used to carry out many important genomic studies about plant, animal, human health and disease^21, 22, 23, 24, 25, 26, 27^. For example, recently, Jin et al. utilized the DNBSEQ platform to enhance our understanding of diseases and phenotypic variations during pregnancy in Asian populations^28^. Since the DNBSEQ and Illumina platforms are both widely used MPS platforms, researchers are concerned about the consistency and interchangeability of genomic variant detection performance between these two platforms. Of which, the performance of SNVs, INDELs and CNVs based on DNBSEQ platforms has been studied and verified to be consistent with those based on Illumina platforms^29, 30, 31^. Specially, in our prior research, we employed five different tools to evaluate the CNV detection capabilities of data derived from the DNBSEQ and Illumina platforms^29^. Our findings indicated that the CNVs identified by both platforms were similar in terms of size, number, sensitivity and precision. Notably, the DNBSEQ platform demonstrated a superior performance in detecting smaller CNVs. However, the comprehensive characteristics of SVs, especially INSs and INVs, identified by DNBSEQ platform remained elusive. To address this, our study embarked on an extensive SV detection analysis, applying 40 different tools to WGS data generated by the DNBSEQ and Illumina platforms for the first time. We meticulously examined their sequencing and genomic attributes to gain a deeper understanding of these variants.

## Results

### Similar SV detection performance between DNBSEQ and Illumina platforms

In our study, we introduced ten WGS datasets of the germline sample NA12878 from public databases^29^ for SV detection, analyzing each dataset with 40 different tools that encompass all five algorithm types as described (Supplementary Data 1, see Supplementary “Methods” for details). Eight datasets were sequenced on two DNBSEQ platforms (BGISEQ-500 and DNBSEQ-G400) with an average depth of 31.43X, and two datasets were sequenced on two Illumina platforms (HiSeq2500 and NovaSeq6000) with an average depth of 30.61X (Supplementary Data 2). Finally, based on DNBSEQ platforms, we detected an average of 2,838 DELs using 32 tools, 1,490 DUPs using 21 tools, 1,117 INSs using 22 tools, 422 INVs using 16 tools, and 2,793 TRAs using eight tools across all eight datasets. These results were very similar to those obtained on the Illumina platforms, including an average of 2,676 DELs, 1,664 DUPs, 737 INSs, 239 INVs, and 2,878 TRAs (Fig.1, Supplementary Data 3).

**Figure 1.**
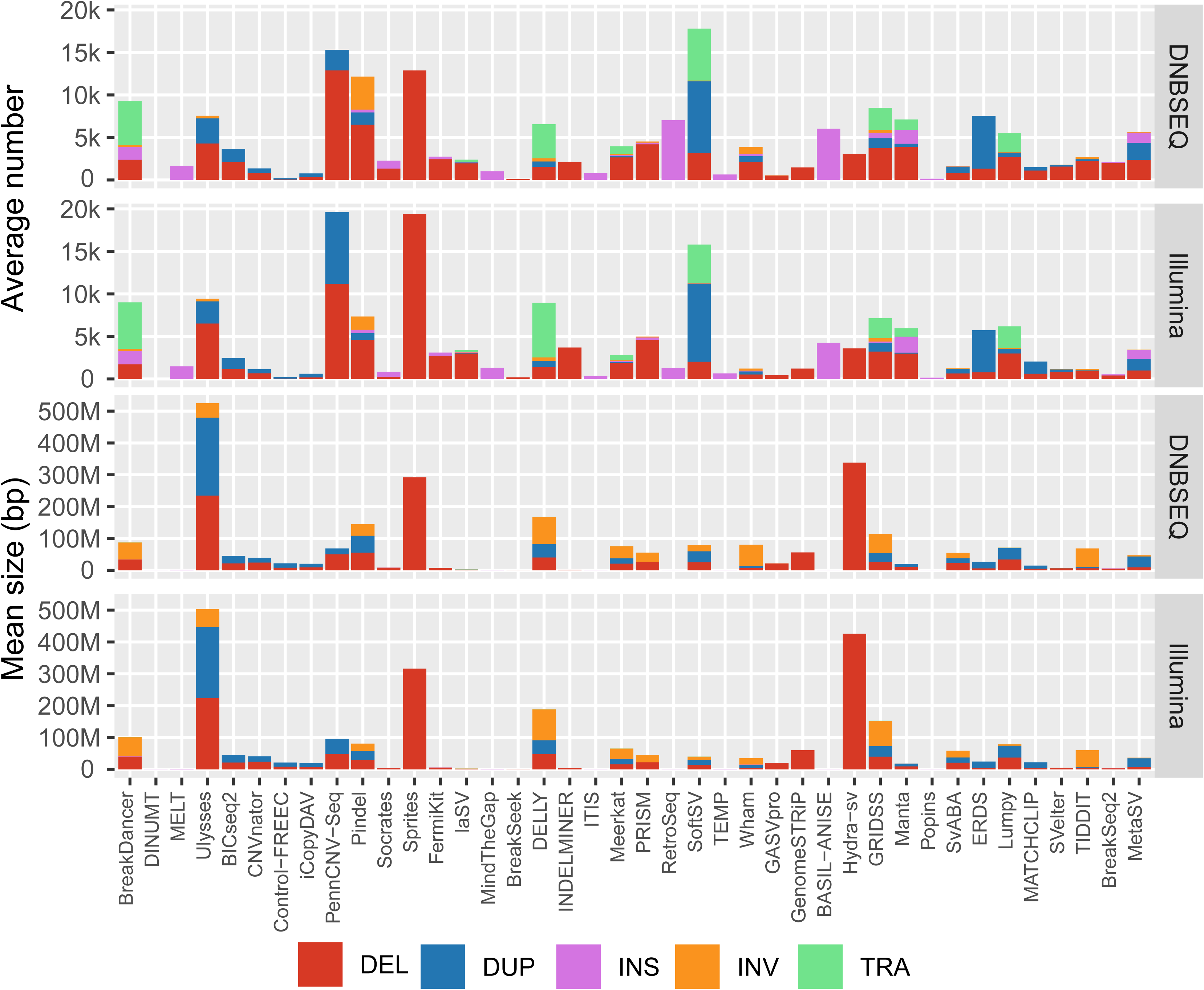
Distribution and Dimensions of SVs. The bar graph illustrates the average number (top panel) and mean size (bottom panel) of SVs identified using the DNBSEQ and Illumina datasets with various analytical tools. SV types are represented by distinct colors: DELs in red, DUPs in blue, INSs in purple, INVs in orange, and TRAs in green. The analytical tools applied for SV detection are displayed along the x-axis. DEL - deletion; DUP - duplication; INS - insertion; INV - inversion; TRA - translocation.

We proceeded to assess the precision and sensitivity of SVs detected by various tools on the DNBSEQ and Illumina platforms. To facilitate a direct comparison with the findings reported by Shunichi et al. in 2019, we adopted their methodologies and benchmarks of the NA12878 sample^13^. In this context, we calculated the precision and sensitivity for DELs, DUPs, INSs, and INVs. TRAs were excluded from the evaluation analysis due to the lack of TRA benchmark of NA12878 and the fact that TRAs are always false positive^32^. The average precision and sensitivity of DELs detected on Illumina datasets were 53.06% and 9.81%, respectively, while 19.86% and 5.52% of DUPs, 44.01% and 2.80% of INSs and 26.79% and 11.06% of INVs were detected, which is consistent with previous report^13^ (Supplementary Fig.1 and Supplementary Data 4). Analogously, the average precision and sensitivity of DELs detected on DNBSEQ datasets were 62.19% and 15.67%, respectively, 23.60% and 6.95% of DUPs, 43.98% and 3.17% of INSs and 25.22% and 11.58% of INVs (Supplementary Fig.2 and Supplementary Data 4). In line with our prior results^29^, the detection of DELs and DUPs on the DNBSEQ platform mirrored the performance observed on the Illumina platform. Meanwhile, the detection of INSs and INVs by the DNBSEQ platform showed a comparable level of performance to those identified using the Illumina platform.

To seek the consistency of genome-wide SV detection between DNBSEQ and Illumina platforms in deep, we compared the number, size, sensitivity and precision of SVs detected by the same tool on these two platforms. We found that both the number and size of various SVs were highly consistent between the DNBSEQ and Illumina platforms (Fig. 2a). Specifically, the consistency of the number and size of DELs were observed with Spearman’s rank correlation coefficients (*rho*) of 0.88 and 0.97 (32 tools), respectively, 0.88 and 0.85 (21 tools) of DUPs, 0.95 and 0.92 (22 tools) of INSs, and 0.96 and 0.88 (16 tools) of INVs (Fig. 2a). Furthermore, the sensitivity and precision of SV detection were also highly consistent between the two platforms, with *rho* values of 0.83 and 0.91 for DELs (Spearman’s rank correlation coefficient), 0.91 and 0.80 for DUPs, 0.96 and 0.97 for INSs, and 0.92 and 0.86 for INVs (Fig. 2b). However, the sensitivity and precision of DELs identified on the DNBSEQ platform (average 15.67% and 62.19%) were found to be marginally higher than those detected on the Illumina platform (9.81% and 53.06%, Fig. 2b, Supplementary Fig.1 and Supplementary Fig.2). Overall, the DNBSEQ and Illumina platforms, both MPS platforms, showed similar SV detection performance, and the SV detection tools developed based on the Illumina platform dataset were also applicable to DNBSEQ platform dataset.

**Figure 2.**
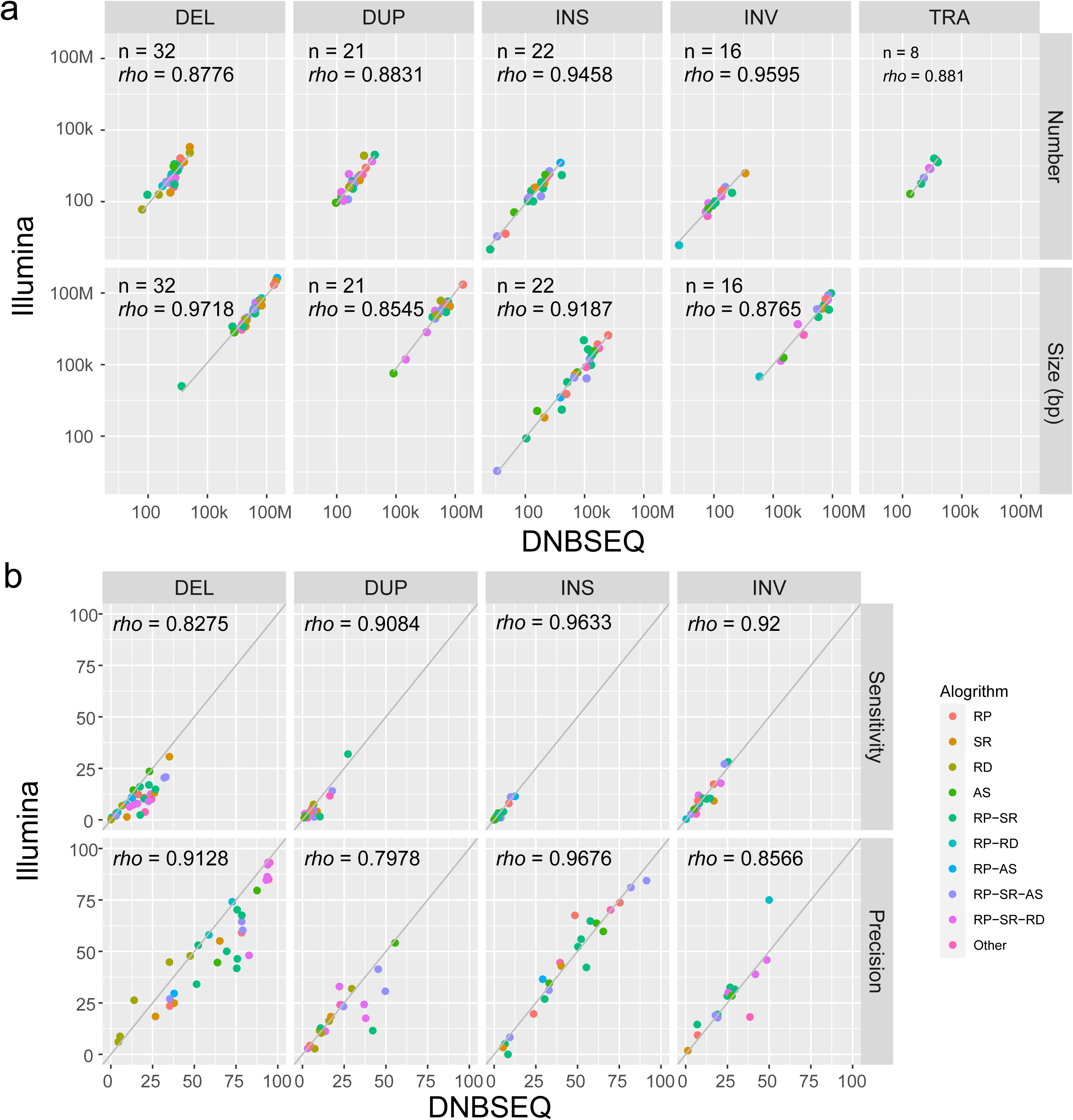
Comparative Analysis of SVs Detected by DNBSEQ and Illumina Platforms. The dot plot presents both the number and size (a), as well as the sensitivity and precision (b), of SVs identified on the DNBSEQ platforms (x-axis) versus those detected on the Illumina platforms (y-axis) using various analytical tools. A total of 40 tools are differentiated by color based on their underlying algorithms. The types of SVs are organized into columns, while the attributes are arranged into rows. The symbol *rho* denotes the correlation coefficient as determined by Spearman’s rank correlation, and n indicates the number of tools capable of detecting a specific type of SV.

### High validation rates of the integrated SV set from DNBSEQ and Illumina platforms

In our study, we built an SV set of DNBSEQ platforms (referred as “DNBSEQ” set) by integrating all SV results of eight DNBSEQ datasets detected by 40 different tools (see Supplementary “Methods” for details). “DNBSEQ” set is consisted of 4,785 SVs, including 3,499 DELs, 630 DUPs, 500 INSs and 156 INVs (Supplementary Fig.3 and Supplementary Data 5). We also integrated SV results of two Illumina datasets and obtained 6,797 SVs, referred as “Illumina” set, including 4,424 DELs, 1,042 DUPs, 1,071 INSs and 260 INVs (Supplementary Data 6).

Presently, three research groups have compiled SVs collections for the NA12878 sample using distinct methodologies. Of which, Marta et al. built a collection of 8,236 SVs from the 1000 Genomes Project (referred as “1KGP” set), which was constructed using high-depth WGS data from the NovaSeq6000^9^. Additionally, Jouni et al. created a collection of 11,089 SVs using the pan-genome approach tailored for short-read data (referred as “Giraffe” set)^33^. Peter et al. employed PacBio RSII long-read data to construct a set of 4,561 SVs for the Human Genome Structural Variation Consortium (referred as “HGSVC” set)^10^. Here, we compared the above-mentioned “DNBSEQ” and “Illumina” sets with “1KGP”, “HGSVC” and “Giraffe” sets. We found that the “DNBSEQ”, “Illumina”, and “1KGP” sets had similar specific SV proportions of 33.38% (1,597/4,785), 51.63% (3,509/6,797) and 30.96% (2,550/8,236), respectively, likely due to their use of short-read MPS data (Supplementary Fig.4). The “Giraffe” set had 66.37% (7,360/11,089) specific SVs, which may be attributed to improvements in pan-genome alignment and analysis methods applied to short-read MPS data, especially for INS detection^33^. The “HGSVC” set had 77.33% (3,527/4,561) specific SVs, likely because it uses long-read sequencing data, highlighting the advantages of long-read data in SV detection^34^.

To enhance the credibility of the “DNBSEQ” and “Illumina” SV sets generated in our research, we employed real-time PCR to confirm the SVs in both SV sets (see Supplementary “Methods” for details). Due to the complexity and variability of the breakpoint regions for DUPs and INVs made it challenging to design reliable primers for these SV types^1, 6^, our validation efforts focused exclusively on DELs and INSs in this study. We randomly selected 17 SVs from “DNBSEQ” and “Illumina” SV sets for real-time PCR validation, including six “DNBSEQ”-specific SVs (three DELs and three INSs), five “Illumina” -specific SVs (three DELs and two INSs), and six shared SVs (three DELs and three INSs, Supplementary Data 7). We designed and synthesized 25 primer pairs targeting the DEL and INS breakpoints identified by Manta software^35^ and conducted real-time PCR assay (Supplementary Fig.5 and Supplementary Fig.6, see Supplementary “Methods” for details). In summary, all 12 SVs from the “DNBSEQ” set were validated via real-time PCR, whereas nine out of 11 SVs within the “Illumina” set were validated (Table 1). In detail, six DELs and six INSs from “DNBSEQ” set were successfully validated, with the exception of one INS breakpoint (chr10:134,865,273). For this INS, only the left breakpoint was validated (Supplementary Fig.6m), while the right breakpoint could not be confirmed (Supplementary Fig.6n). In contrast, within the “Illumina” set, four DELs and five INSs were confirmed, while two DELs failed to be validated by real-time PCR. Moreover, all six shared SVs (three DELs and three INSs) showed a validation rate of 100%. Given the high validation rates of both the “DNBSEQ” and “Illumina” SV sets, we further analyzed their genomic characterizes without additional modifications.

### Genomic characterizing the SVs of the “DNBSEQ” and “Illumina” sets

Analyzing the genomic characteristics of SVs, such as repetitive DNA composition and GC content, provides valuable insights into the origins, mechanisms, and functional impacts of SVs, and even are crucial for disease research, evolutionary biology, and understanding genomic functions^36^. After evaluating the consistency of SVs from the DNBSEQ and Illumina datasets using various tools, we further analyzed and compared the genomic characteristics of SVs from both sets to explore their overall consistency. Firstly, we analyzed the repetitive DNA components and sequences of SVs in our “DNBSEQ” and “Illumina” sets. We found the size distribution of SV was similar between these two sets, with both showing mobile element signatures of Alu (∼300 bp) and LINE1 (L1, ∼6 kb, Fig.3a), which is consistent with previous report^9, 34^. We also annotated the SVs to repeat regions and found that the majority were located in repeat regions (Fig.3b). Specifically, 24.20% of SVs in the “DNBSEQ” set and 38.88% in the “Illumina” set were associated with tandem repeats (detected by Tandem Repeats Finder, TRF). This was followed by short tandem repeats (STR) at 27.61% and 24.51% in “DNBSEQ” and “Illumina”, respectively, Alu elements at 26.04% and 16.57%, and L1 elements at 14.69% and 12.89%. Only a small percentage of SVs were not located in repeat regions, including 0.40% in “DNBSEQ” and 0.38% in “Illumina”. These findings suggest that SVs are clustered in repeat regions on the human genome^34^, and that both “DNBSEQ” and “Illumina” platforms are capable of detecting SVs in these regions.

**Figure 3.**
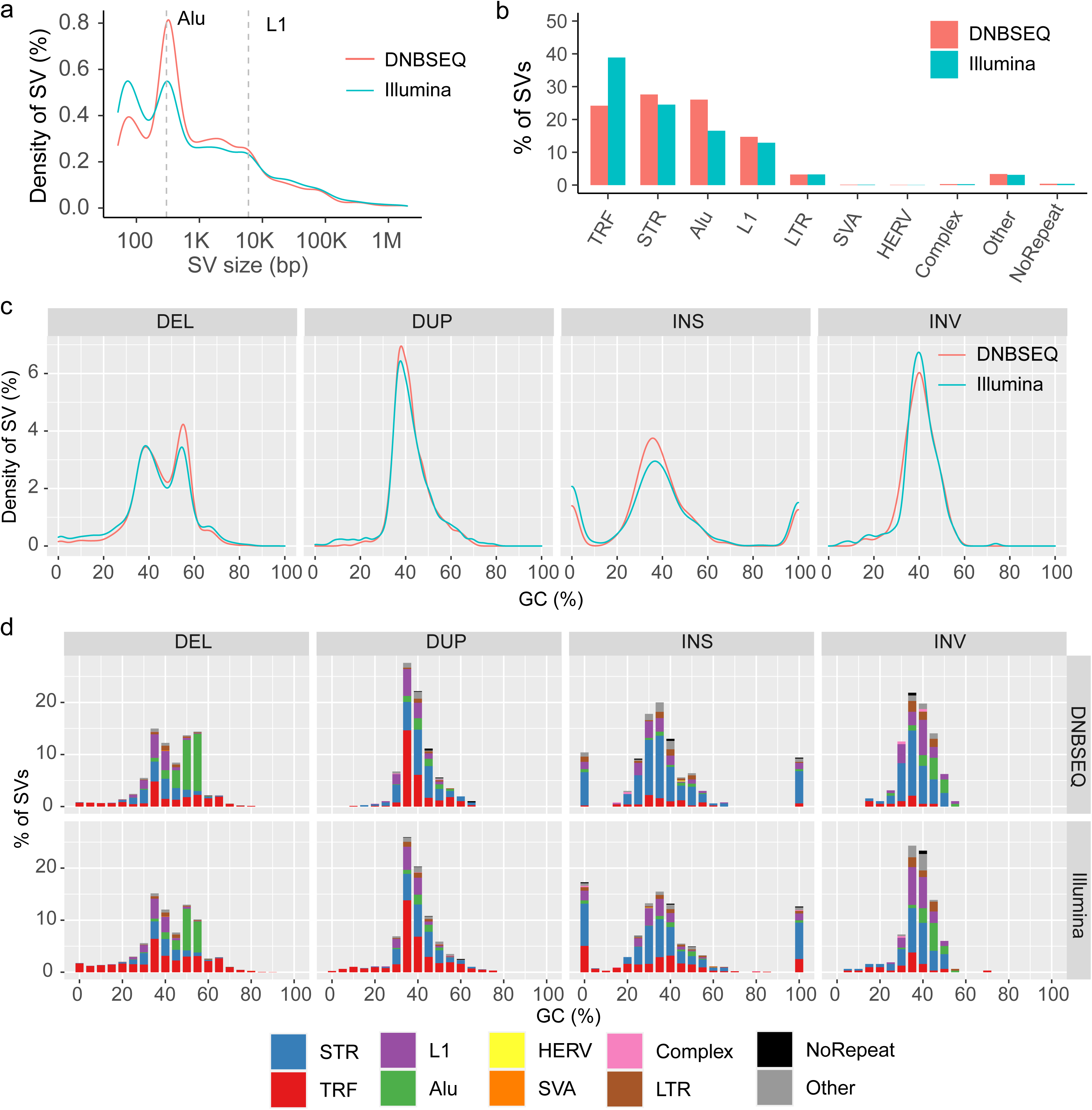
Characteristics of DNBSEQ and Illumina SV Sets. (a) The size distribution density for DNBSEQ and Illumina SVs reveals a peak at 300 bp corresponding to Alu elements and a 6 kb peak associated with L1 elements. (b) The bar chart illustrates the percentage of DNBSEQ and Illumina SVs (y-axis) that are annotated within specific repetitive regions (x-axis). (c) The line density plot depicts the distribution of GC content across each type of SV (column), with the y-axis representing the proportion of SVs at a given GC content level (x-axis). (d) The bar chart details the repetitive components within SVs of varying GC content for each SV category (column), showing the percentage of repetitive regions (y-axis) to which SVs of particular GC contents (x-axis) are annotated. The GC content is categorized into bins at intervals of 5% GC. The DNBSEQ and Illumina SV sets are distinguished by color: DNBSEQ set, red; Illumina set, green. Different colors denote the repetitive regions: STR, short tandem repeats, blue; TRF, tandem repeats detected by Tandem Repeats Finder, red; L1, LINE1, purple; Alu, green; HERV, human endogenous retroviruses, yellow; SVA, orange; Complex, low complexity, pink; LTR, long tandem repeats, brown; Other, gray; NoRepeat, black.

We also analyzed the GC composition of SVs in the “DNBSEQ” and “Illumina” sets to better understand their sequence components. We found that, regardless of the SV set, the GC distribution of DUPs and INVs were close to the reference distribution, with enrichment in the 40-50% GC content range, while DELs and INSs exhibited a different pattern (Fig.3c). DELs in both “DNBSEQ” and “Illumina” sets exhibited a bimodal GC distribution with peaks at 40% and 55%. Specifically, 34.55% (1,209/3,499) of DELs in the “DNBSEQ” set and 28.28% (1,251/6,797) of DELs in the “Illumina” set had a GC content in the 50-60% range, corresponding to the 55% peak (Fig.3d). Among these DELs with 50-60% GC content, 73.45% (888/1,209) in the “DNBSEQ” set and 61.47% (760/1,251) in the “Illumina” set were associated with Alu sequences (Fig.3d). In contrast to DELs, INSs in both the DNBSEQ and Illumina sets exhibited GC content distributions with peaks at 0%, 40%, and 100%. Among INSs with 0% and 100% GC content, 63.64% in the “DNBSEQ” set and 50.63% in the “Illumina” set were STR sequences, suggesting that this enrichment in the extreme GC content of INSs was primarily driven by STRs (Fig.3d). These findings suggest that the GC content of SVs varies depending on the type of SV and the repeat elements involved. Despite these differences, the GC content patterns of SVs were consistent between the “DNBSEQ” and “Illumina” sets.

Previous study had reported that SVs exhibit non-random distribution patterns in the genome, with enrichment in repeat regions and a notable bias towards chromosomal ends^34^. Our analysis confirmed that both the “DNBSEQ” and “Illumina” SVs exhibit repeat region enrichment consistent with this report (Fig.4a). Furthermore, we analyzed the chromosomal location biases of the “DNBSEQ” and “Illumina” SVs. We observed a 1.26-fold (p = 4.726e-06, z-score = 4.58, permutation test) enrichment of “DNBSEQ” SVs and 1.58-fold (p < 2.2e-16, z-score = 8.31, permutation test) enrichment of “Illumina” SVs within 5 Mbp of telomere, respectively, which is consistent with the findings of earlier study^34^. However, we also found that 15.01% of “DNBSEQ” SVs (718/4,785) and 15.99% of “Illumina” SVs (1,087/6,797) were located within 5 Mbp of centromere, presenting a 2.03-fold (p < 2.2e-16, z-score = 15.37, permutation test) and 2.34-fold (p < 2.2e-16, z-score = 16.56, permutation test) enrichment, respectively (Supplementary Fig.7). Since SVs were known to be clustered^34^, we further identified hotspots in the “DNBSEQ” and “Illumina” sets and analyzed the chromosomal distribution of these hotspots (Fig.4a). We identified 26 SV hotspots in the “DNBSEQ” set and 51 SV hotspots in the “Illumina” set (Supplementary Data 8 and Supplementary Data 9). Of these, 30.77% (8/26) hotspots in the “DNBSEQ” set and 31.37% (16/51) hotspots in the “Illumina” set were previously reported in research using long-read datasets^10^. As expected, these hotspots in both the “DNBSEQ” and “Illumina” sets were mostly located near centromeres or telomeres.

Given that it is challenging to clone and sequence regions with GC-bias^34^, we sought to assess the performance of “DNBSEQ” and “Illumina” SVs in these difficult genomic regions. To achieve this, we classified the “DNBSEQ” and “Illumina” SVs according to the difficult- and easy-to-sequence regions defined by the Genome in a Bottle (GIAB) Consortium^37^. We found that 59.16% of “DNBSEQ” SVs (2,831/4,785) and 65.18% of “Illumina” SVs (4,430/6,797) were located in difficult-to-sequence regions, even though these regions only make up 18.00% of the human genome (Fig.4b). In detail, 65.36% of DELs (2,287/3,499) and 72.70% of DUPs (458/639) in the “DNBSEQ” set were enriched in difficult-to-sequence regions, whereas only 9.40% of INSs (47/500) and 25.00% of INVs (39/156) were found in these regions (Fig.4b). Similarly, in the Illumina set, 73.73% of DELs (3,262/4,424) and 85.12% of DUPs (887/1,042) were located in difficult-to-sequence regions, compared to 18.02% of INSs (193/1,071) and 33.85% of INVs (88/260). These findings suggest that SVs are more likely to occur in specific genomic regions, including repeats and difficult-to-sequence regions, and exhibit GC-biases in their chromosomal locations. Additionally, we annotated SVs to functional region of human genes to gain a better understanding of their impact on functional regions. We found that 62.28% of “DNBSEQ” SVs (2,980/4,785) and 62.70% of “Illumina” SVs (4,262/6,797) were located in intergenic regions, followed by 35.42% of “DNBSEQ” SVs (1,695/4,785) and 34.38% of “Illumina” SVs (2,337/6,797) in intronic regions (Fig.4c). Only 2.03% of “DNBSEQ” SVs and 2.91% of “Illumina” SVs intersected with functional elements such as exons (26 “DNBSEQ” SVs and 40 “Illumina” SVs), promoters (25 and 46), and UTRs (8 and 22). Interestingly, 81.44% of “DNBSEQ” SVs (3,897/4,785) and 81.48% of “Illumina” SVs (5,538/6,797) were located within 0.25 Mb of transcription start site (TSS, Supplementary Fig.8). Our analysis revealed that SVs in both the “DNBSEQ” and “Illumina” sets are predominantly located in intergenic regions and depleted in functional regions of genes, such as exons and introns. In conclusion, we further validated the consistency of SV sets from both DNBSEQ and Illumina platforms across multiple genomic characteristics, including repetitive regions, GC distribution, difficult-to-sequence regions, and gene features. This consistency underscores the robustness of our comparative analysis and highlights the value of both platforms in understanding the genomic context of SVs.

### Resource consumption

We recorded the time and memory of SV detection on ∼30X WGS datasets using various tools (Supplementary Data 10). We found that Sprites (mean = 229.77 h and mean = 439.68 h on DNBSEQ and Illumina datasets, respectively, Supplementary Fig.9), Pindel^38^ (216.68 h and 163.32 h), MindTheGap^39^ (166.92 h and 190.65 h), and laSV^40^ (114.69 h and 121.17 h) had large time consumption, while laSV (mean = 150.91 GB and 143.47 GB on DNBSEQ and Illumina datasets, respectively), FermiKit^41^ (73.52 GB and 54.42 GB), and MindTheGap (33.61 GB and 39.93 GB) had large memory consumption. This differences in time and memory consumption were related to the input format and algorithms of tools. However, we found high consistency of time consumption and memory consumption between the DNBSEQ and Illumina datasets using the same tools under similar ∼30X data size (*rho* = 0.97 for time consumption and 0.95 for memory consumption, Spearman’s rank correlation coefficient). These findings suggest that the choice of SV detection tool may depend on the specific needs of the study, such as the desired balance between time and memory consumption.

## Discussion

Although SV detection on the Illumina platforms has increasingly demonstrated the importance of SVs, there is insufficient information regarding the performance of SV detection on another widely used MPS platform: the DNBSEQ platforms. In this study, we detected and characterized SVs, including DELs, DUPs, INSs and INVs, using DNBSEQ and Illumina datasets with 40 tools for the first time. Overall, our systematic analysis demonstrated that the DNBSEQ platform exhibits performance in SV detection that is consistent with the Illumina platform across various aspects, including the number, size, precision, and sensitivity of detected SVs, as well as their composition in repeats, genomic element distribution, and genomic localization.

Various tools have been designed to detect SVs using short-read signatures based on dataset sequenced on Illumina platforms. These tools have shown varying levels of precision and sensitivity for different SV types. This study demonstrated that SV detection tools developed for Illumina dataset are also compatible with the DNBSEQ dataset, as the results and performance of SV detection were consistent between the DNBSEQ and Illumina datasets using the same tool. However, we also observed notable performance differences between different tools, regardless of whether they were applied to DNBSEQ or Illumina datasets (Supplementary Data 4 and 11). For example, Manta, GRIDSS, SoftSV, and MetaSV showed higher precision and sensitivity in detecting deletions and duplications, while MELT was a good choice for insertion detection. TIDDIT, DELLY, and GRIDSS were good at detecting inversions. In contrast, using the same given tool for SV detection, we achieved an average concordance rate of 55.40% for DNBSEQ datasets and 40.29% for Illumina datasets. These results not only confirm the consistency between the DNBSEQ and Illumina platforms but also highlight the importance and necessity of carefully selecting SV detection software, regardless of the data platform used.

Recent advancements in sequencing and data analysis technologies have significantly enhanced our detection and understanding of SVs. Long-read sequencing technology can fully sequence large DNA fragments (>10 kb), providing continuous sequence that spans entire SVs^34^. For short-read dataset, pangenomic technology, particularly Giraffe, reduces reference allele bias and improves SV detection performance by mapping reads to a haplotype-resolved graph that includes references from thousands of human genomes^33^. In this study, we demonstrated the ability to detect SVs on short-read WGS data using DNBSEQ platforms and characterized the genomic features of these SVs. Our results showed a high consistent ratio of NA12878 SVs between “DNBSEQ”, “Illumina”, and “1KGP” sets, all of which were sequenced on short-read WGS data based on MPS platforms. For example, 73.34%, 53.05%, and 78.67% of DELs in the “DNBSEQ”, “Illumina”, and “1KGP” sets were shared, respectively (Supplementary Fig.4). However, these three SV sets exhibit notable differences compared to the “HGSVC” set, which uses long-read data, and the “Giraffe” set, which employs pangenomic technology. For instance, 76.31% of DELs and 77.92% of INSs in the “HGSVC” set, and 61.07% of DELs and 70.53% of INSs in the “Giraffe” set, could not be detected in any of “DNBSEQ”, “Illumina”, or “1KGP” sets. This result highlights the limitation of SVs detection, especially INSs detection, by normally short-read datasets, which is consistent with previous research^12, 33^. The distributions of SVs on short- and long-read platforms were also found to be inconsistent. Most of the SVs on long-read platforms were concentrated within 5 Mbp of the telomere^34^, while the SVs on short-read platforms were enriched within 5 Mbp of both the telomere and centromere (Fig.4; Supplementary Fig.7). These results confirm the complexity of SV detection and illustrate the stability and limitation of SV detection based on short-read MPS platforms. In our future work, we will continue to explore the performance of SVs detection in the DNBSEQ datasets combined with pangenomic technology, as well as analyze the SV detection performance of long-read platforms, such as PacBio^42^, Oxford Nanopore^43^ and CycloneSEQ^44^.

**Figure 4.**
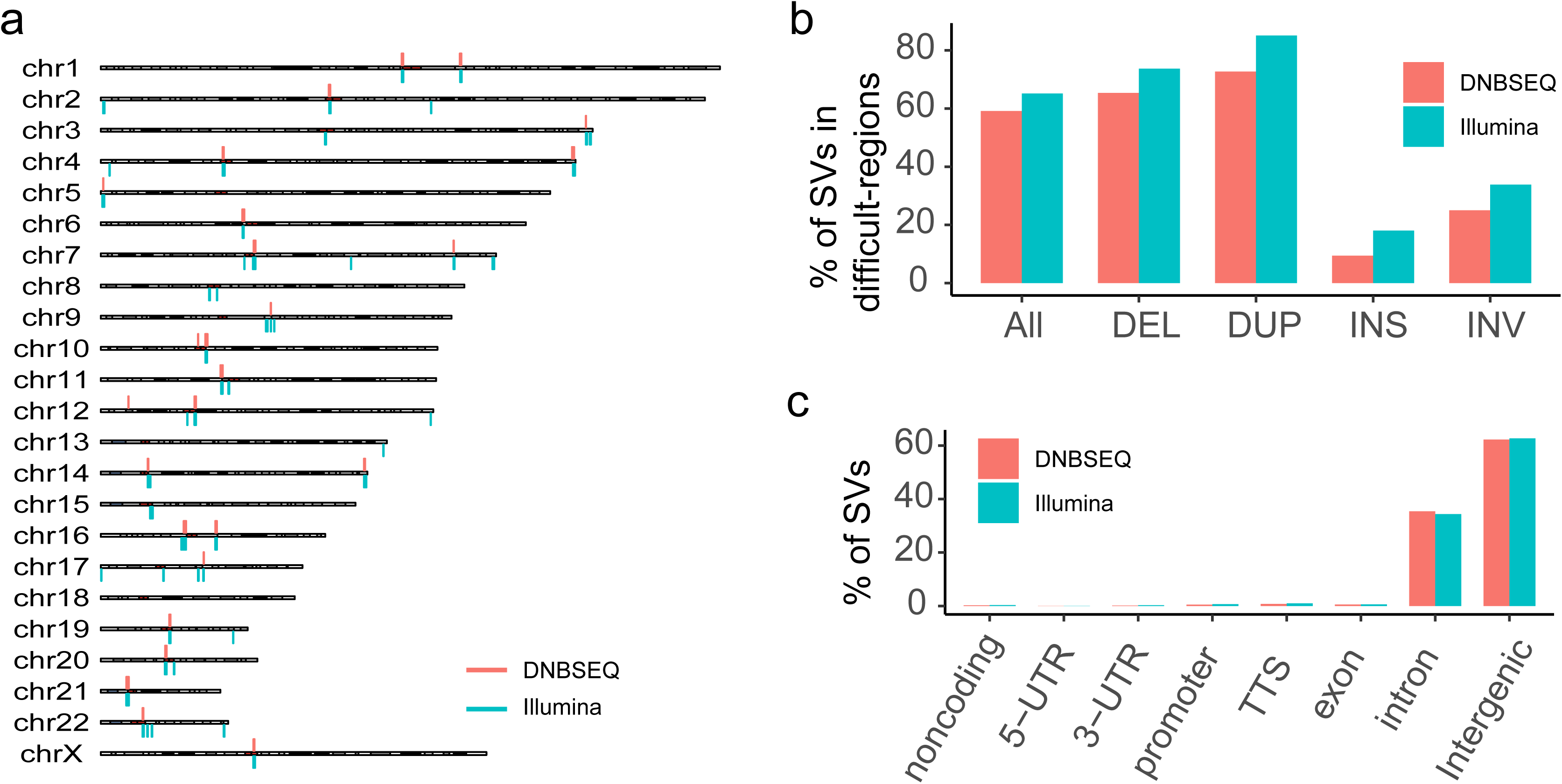
Distribution of SVs. (a) The bar chart displays the hotspots of DNBSEQ and Illumina SV sets on human genome. The gray and white bands on ideogram indicate different genomic regions, and the red bands represent centromeres. (b) The bar chart indicates the percentage of DNBSEQ and Illumina SVs located within genomic regions that are typically difficult to sequence or analyze. The proportions are shown on the y-axis, while the x-axis categorizes the different types of SVs. (c) The bar chart illustrates the distribution of DNBSEQ and Illumina SVs across various gene regions, with the y-axis indicating the proportion of SVs mapped to specific gene regions on the x-axis. The DNBSEQ and Illumina SV sets are differentiated by color: DNBSEQ, DNBSEQ SV set, red; Illumina, Illumina SV set, blue.

In conclusion, we systematically analyzed the performance and characteristics of germline SVs detected in WGS datasets sequenced on the DNBSEQ platform. By evaluating the performance of SV detection with the same tool and integrating the results of all tools to assess the genomic characteristics of SV sets, our study demonstrated the consistency of SV detection between the DNBSEQ and Illumina platforms. Furthermore, we provided a benchmark reference for future SV detection based on the DNBSEQ platforms.

## Methods

### Sequencing data resources

Ten WGS datasets of NA12878 were utilized for SV detection, which were publicly available and analyzed in our previous article^29^. All ten WGS datasets of NA12878 were processed as previously described, with an average coverage of approximately 30X, mapping rate of over 99%, and genome coverage of over 99%.

### Public SV sets of NA12878

Two public SV sets of NA12878 were downloaded and compared: 8,469 SVs detected using NovaSeq6000 by Marta et al. (referred to as “1KGP” ^9^, 4,718 SVs detected using PacBio RSII by Peter et al. (referred to as “HGSVC”)^34^ and 11,320 SVs (genotype quality ≥ 60) detected using Giraffe by Jouni et al. (referred to as “Giraffe”)^33^. All three SV sets were converted to BED format and lifted over from GRCh38 to hg19 using the LiftOver tool (UCSC)^45^. This resulted in 8,236, 4,561 and 11,089 SVs of “1KGP”, “HGSVC” and “Giraffe” sets, respectively.

### SV detection

Forty tools were utilized for SV detection on the ten WGS datasets of NA12878, respectively. There 40 tools were carefully selected based on the following main factors: ability to work on individual WGS data, ability to detect SVs on real data, suitability for MPS short-read data and ease of accessibility. The SV calling were processed according to the approach described by Shunichi et al.^13^. SVs meeting the following criteria were filtered out: (1) the size of DEL, DUP and INV > 2M bp or < 50 bp, (2) the number of reads supporting the called SV (RSS) < 3, (3) not located on autosomal or chrX chromosomes, or (4) overlapping a gap in the reference genome.

### SV evaluation

A reference dataset of SV in NA12878, as described in Shunichi et al.^13^ was downloaded to evaluate SV performance (https://github.com/stat-lab/EvalSVcallers/blob/master/Ref_SV/NA12878_DGV-2016_LR-assembly.vcf). The dataset contained 9,241 DELs, 2,611 DUPs, 13,669 INSs and 291 INVs. The evaluation of SVs was performed using a script from https://github.com/stat-lab/EvalSVcallers based on a benchmark. DEL, DUP or INS was judged as true positive if it had a reciprocally overlap (RO) of ≥ 50% with the reference DEL, DUP or INS, respectively. INS was judged as true positive if the breakpoints of the called INS were located within +/− 200bp of those of the reference INS. The precision, sensitivity and F1-score were calculated using the following equations:

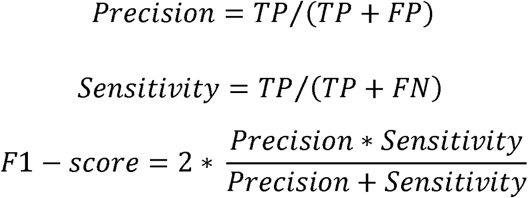

where TP is the true positive of SV, FP is the false positive, and FN is the false negative.

### SV integration

We created a union set of SVs by integrating all SV results for each SV variant type (i.e., DELs were integrated with other DELs, and the same for DUPs, INSs and INVs). We integrated the SV set by merging pairwise comparison results sequentially in the order of the sample list in Supplementary Data 2. Specifically, the result of pairwise comparison between the first two datasets was compared with the third dataset, and so on, until all data were used. DELs, DUPs and INVs were excluded if they had a ≥ 50% RO with other DELs, DUPs or INVs in pairwise comparison, but INSs were excluded if they were located within 200bp of other INSs. The common SVs detected in all datasets sequenced on the same platform (i.e., DNBSEQ or Illumina platforms) using the same tool were defined as candidate SV set of that tool, and the candidate SVs detected by two or more tools were merged as the SV set of the platform. Finally, we obtained 4,785 SVs on DNBSEQ platforms (Supplementary Data 5) and 6,797 SVs on Illumina platforms (Supplementary Data 6), respectively. The low number of SVs in the DNBSEQ set is primarily due to the inherently low consistency ratio between SVs detected on different datasets using the same tool (mean = 48.97%, range 0.09% - 96.93%, Supplementary Fig.10). Therefore, the DNBSEQ set from DNBSEQ platforms with more integrated datasets has a lower number of SVs.

### SV validation with real-time PCR

We randomly selected DELs and INSs from the DNBSEQ and Illumina datasets for validation, identifying the sequences of these SVs using Manta. However, DUPs and INVs were not included in the validation process due to the unavailability of their breakpoint sequences necessary for primer design. Primer sequences were crafted using Primer Premier 6.0 and synthesized by BGI-Write (Supplementary Data 7). Human genomic DNA of NA12878 was purchased at the Coriell Institute. Real-time PCR assays were conducted using SYBR Green, with all procedures performed on the StepOne Real-Time PCR System (Applied Biosystems) in accordance with the manufacturer’s protocol. For each breakpoint, seven real-time PCR reactions were executed as per the manufacturer’s guidelines. This included triplicate reactions for the target primers, a no-template control to ensure specificity, and two positive controls: one using a standard sample and the other employing the GAPDH gene as an internal reference.

### Identification of SV hotspots

The midpoint of the SV region was extracted and used to identify hotspot on hg19 genome. The midpoint sites were transformed using the ‘makeGRangesFromDataFrame’ function from the GenomicRanges package(v1.24.1) and then submitted to the ‘hotspotter’ function from the primatR package (with parameters: bw = 200000, num.trial = 1000). The hotspots were displayed on hg19 genome using the karyoploteR package (v1.20.0).

### Annotation of difficult regions

The difficult regions of hg19 were downloaded from the GIAB (https://ftp-trace.ncbi.nlm.nih.gov/giab/ftp/release/genome-stratifications/v3.1/GRCh37/Union/GRCh37_alldifficultregions.bed.gz), which including 5,427,803 regions spanning 557.28 Mbp. We classified SVs as being in the difficult region if they were located in any difficult region with >50% size overlap, and the remaining SVs were defined as being in the easy region.

### Annotation of genes

The SVs in the DNBSEQ and Illumina sets were annotated to hg19 genes using HOMER annotatePeaks.pl (v4.11). We extracted the gene annotation and distance to TSS from the output of HOMER annotatePeaks.pl.

### Annotation of repeat regions

The TRF and rmsk regions were downloaded from the UCSC Genome Browser (https://hgdownload.soe.ucsc.edu/goldenPath/hg19/database/), while the SVA regions were obtained from MELT (v2.2.2.2). DELs, DUPs and INVs were annotated to TRF, rmsk and SVA regions, respectively, when >50% of the SV size was located in a repeat region using BEDTools (v2.30.0). INSs were annotated if they were within 50bp of any repeat regions. In case where a single SV was annotated to multiple repeat regions, we selected the superior annotation based on the following priorities: TRF > STR > Alu > L1 > SVA > HERV > LTR > Low Complexity > Other Repeats.

## Availability of data and materials

All data analyzed in this study are included in previous published article and its supplementary files.

## Abbreviations

*SNV*: single-nucleotide variant
*INDEL*: insertion and deletion
*WGS*: whole-genome sequencing
*DEL*: deletion
*DUP*: duplication
*INS*: insertion
*INV*: inversion
*STR*: short tandem repeat
*MPS*: massively parallel sequencing

## Acknowledgements

This work was supported by the National Key Research and Development Program of China (2021YFF1200105).

## Author information

### Contributions

J.R., L.P. and F.C. designed the project. H.L. performed the experiment. J.R., H.L., D.A. and X.L. performed the data analysis and involved in the data curation. J.R. and H.L. wrote the manuscript. J.R., L.P. and F.C. revised the manuscript. All authors reviewed and approved the final manuscript.

## Ethics declarations

### Ethics approval and consent to participate

Not applicable.

### Consent for publication

Not applicable.

## References

1. Alkan C, Coe BP, Eichler EE. Genome structural variation discovery and genotyping. Nat Rev Genet 12, 363–376 (2011).

2. Abyzov A, et al. Analysis of deletion breakpoints from 1,092 humans reveals details of mutation mechanisms. Nat Commun 6, 7256 (2015).

3. Feuk L, Carson AR, Scherer SW. Structural variation in the human genome. Nat Rev Genet 7, 85–97 (2006).

4. Stankiewicz P, Lupski JR. Structural variation in the human genome and its role in disease. Annu Rev Med 61, 437–455 (2010).

5. Sudmant PH, et al. Global diversity, population stratification, and selection of human copy-number variation. Science 349, aab3761 (2015).

6. Sudmant PH, et al. An integrated map of structural variation in 2,504 human genomes. Nature 526, 75–81 (2015).

7. Weischenfeldt J, Symmons O, Spitz F, Korbel JO. Phenotypic impact of genomic structural variation: insights from and for human disease. Nat Rev Genet 14, 125–138 (2013).

8. Collins RL, et al. A structural variation reference for medical and population genetics. Nature 581, 444–451 (2020).

9. Byrska-Bishop M, et al. High-coverage whole-genome sequencing of the expanded 1000 Genomes Project cohort including 602 trios. Cell 185, 3426–3440 e3419 (2022).

10. Ebert P, et al. Haplotype-resolved diverse human genomes and integrated analysis of structural variation. Science 372, (2021).

11. Collins RL, et al. A cross-disorder dosage sensitivity map of the human genome. Cell 185, 3041–3055 e3025 (2022).

12. Zhao X, et al. Expectations and blind spots for structural variation detection from long-read assemblies and short-read genome sequencing technologies. Am J Hum Genet 108, 919–928 (2021).

13. Kosugi S, Momozawa Y, Liu X, Terao C, Kubo M, Kamatani Y. Comprehensive evaluation of structural variation detection algorithms for whole genome sequencing. Genome Biol 20, 117 (2019).

14. Abyzov A, Urban AE, Snyder M, Gerstein M. CNVnator: an approach to discover, genotype, and characterize typical and atypical CNVs from family and population genome sequencing. Genome Res 21, 974–984 (2011).

15. Holtgrewe M, Kuchenbecker L, Reinert K. Methods for the detection and assembly of novel sequence in high-throughput sequencing data. Bioinformatics 31, 1904–1912 (2015).

16. Zhang Z, et al. Sprites: detection of deletions from sequencing data by re-aligning split reads. Bioinformatics 32, 1788–1796 (2016).

17. Mahmoud M, Gobet N, Cruz-Davalos DI, Mounier N, Dessimoz C, Sedlazeck FJ. Structural variant calling: the long and the short of it. Genome Biol 20, 246 (2019).

18. Chaisson MJP, et al. Multi-platform discovery of haplotype-resolved structural variation in human genomes. Nat Commun 10, 1784 (2019).

19. Zhao M, Wang Q, Wang Q, Jia P, Zhao Z. Computational tools for copy number variation (CNV) detection using next-generation sequencing data: features and perspectives. BMC Bioinformatics 14 Suppl 11, S1 (2013).

20. Drmanac R, et al. Human genome sequencing using unchained base reads on self-assembling DNA nanoarrays. Science 327, 78–81 (2010).

21. Zhou P, et al. A pneumonia outbreak associated with a new coronavirus of probable bat origin. Nature 579, 270–273 (2020).

22. Wang K, et al. African lungfish genome sheds light on the vertebrate water-to-land transition. Cell 184, 1362–1376 e1318 (2021).

23. Liu S, et al. Utilizing non-invasive prenatal test sequencing data for human genetic investigation. Cell Genomics 4, (2024).

24. Zhu H, et al. Novel insights into the genetic architecture of pregnancy glycemic traits from 14,744 Chinese maternities. Cell Genomics 4, (2024).

25. Xiao H, et al. Genetic analyses of 104 phenotypes in 20,900 Chinese pregnant women reveal pregnancy-specific discoveries. Cell Genomics 4, (2024).

26. Liu S, et al. Genome-wide association study of maternal plasma metabolites during pregnancy. Cell Genomics 4, (2024).

27. Guo J, et al. Phenome-wide association study in 25,639 pregnant Chinese women reveals loci associated with maternal comorbidities and child health. Cell Genomics 4, (2024).

28. Jin X, et al. Advances in using non-invasive prenatal testing to study genomics related to maternity. Cell Genomics 4, (2024).

29. Rao J, et al. Performance of copy number variants detection based on whole-genome sequencing by DNBSEQ platforms. BMC Bioinformatics 21, 518 (2020).

30. Huang J, et al. A reference human genome dataset of the BGISEQ-500 sequencer. Gigascience 6, 1–9 (2017).

31. Galli A, et al. Germline and somatic variant identification using BGISEQ-500 and HiSeq X Ten whole genome sequencing. Plos One 13, (2018).

32. Sedlazeck FJ, et al. Accurate detection of complex structural variations using single-molecule sequencing. Nat Methods 15, 461–468 (2018).

33. Sirén J, et al. Pangenomics enables genotyping of known structural variants in 5202 diverse genomes. Science 374, (2021).

34. Audano PA, et al. Characterizing the Major Structural Variant Alleles of the Human Genome. Cell 176, 663–675 e619 (2019).

35. Chen X, et al. Manta: rapid detection of structural variants and indels for germline and cancer sequencing applications. Bioinformatics 32, 1220-1222 (2016).

36. Ho SS, Urban AE, Mills RE. Structural variation in the sequencing era. Nat Rev Genet 21, 171–189 (2020).

37. Krusche P, et al. Best practices for benchmarking germline small-variant calls in human genomes. Nat Biotechnol 37, 555–560 (2019).

38. Ye K, Schulz MH, Long Q, Apweiler R, Ning Z. Pindel: a pattern growth approach to detect break points of large deletions and medium sized insertions from paired-end short reads. Bioinformatics 25, 2865–2871 (2009).

39. Rizk G, Gouin A, Chikhi R, Lemaitre C. MindTheGap: integrated detection and assembly of short and long insertions. Bioinformatics 30, 3451–3457 (2014).

40. Zhuang J, Weng Z. Local sequence assembly reveals a high-resolution profile of somatic structural variations in 97 cancer genomes. Nucleic Acids Res 43, 8146–8156 (2015).

41. Li H. FermiKit: assembly-based variant calling for Illumina resequencing data. Bioinformatics 31, 3694–3696 (2015).

42. Eid J, et al. Real-time DNA sequencing from single polymerase molecules. Science 323, 133–138 (2009).

43. Loman NJ, Quick J, Simpson JT. A complete bacterial genome assembled de novo using only nanopore sequencing data. Nature Methods 12, 733–735 (2015).

44. Zhang J-Y, et al. A single-molecule nanopore sequencing platform. bioRxiv, 2024.2008.2019.608720 (2024).

45. Hinrichs AS. The UCSC Genome Browser Database: update 2006. Nucleic Acids Research 34, D590–D598 (2006).

