## Supplementary Fig for "Performance evaluation of structural variation detection using DNBSEQ whole-genome sequencing"

*Rao et al.*

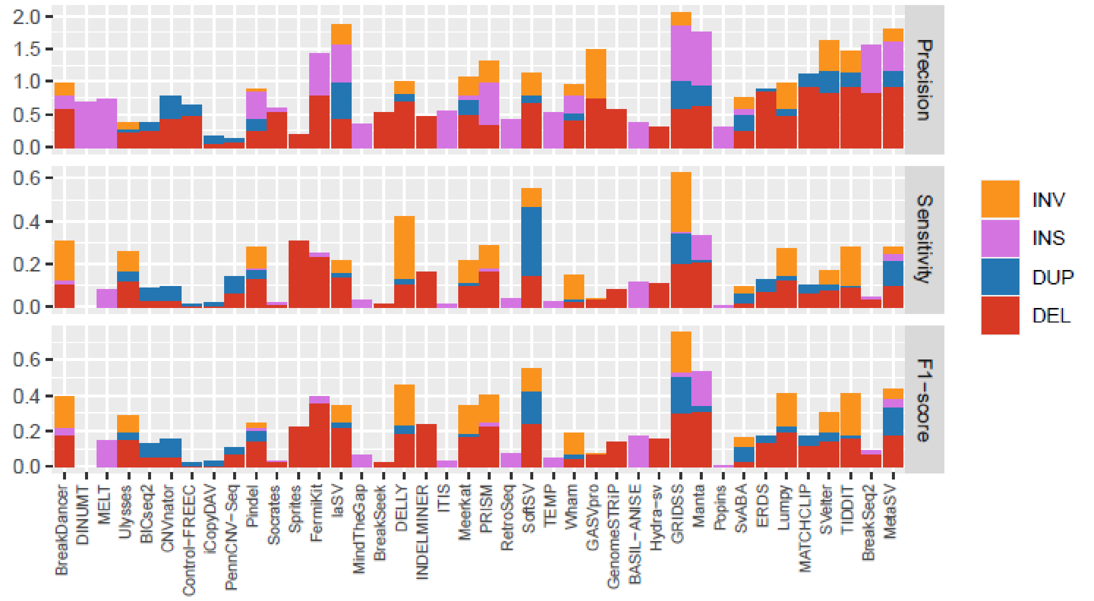


**Supplementary Figure 1. Precision, sensitivity and F1-score of SVs detected on DNBSEQ datasets.** Histogram shows the mean precision (**top**), mean sensitivity (**middle**) and mean F1-score (**bottom**) of SVs identified in DNBSEQ datasets using various tools, as indicated on the x-axis. The performance metrics for each type of SV are color-coded: DEL, deletion, red; DUP, duplication, blue; INS, insertion, purple; INV, inversion, orange.

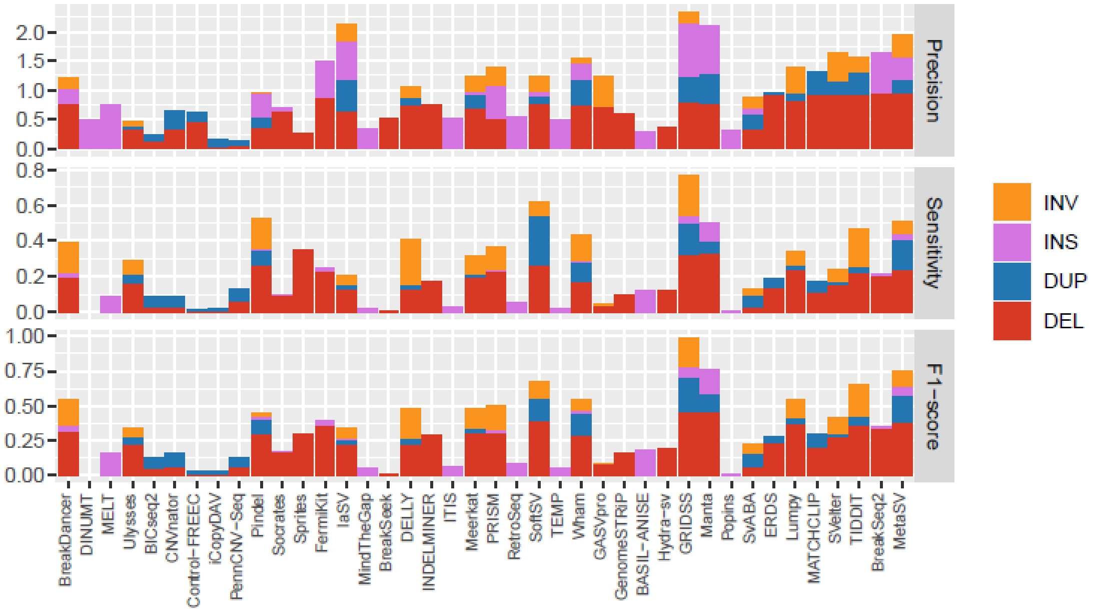


**Supplementary Figure 2. Precision, sensitivity and F1-score of SVs detected on Illumina datasets.** Histogram shows the mean precision (**top**), mean sensitivity (**middle**) and mean F1-score (**bottom**) of SVs identified in Illumina datasets using various tools, as indicated on the x-axis. The performance metrics for each type of SV are color-coded: DEL, deletion, red; DUP, duplication, blue; INS, insertion, purple; INV, inversion, orange.

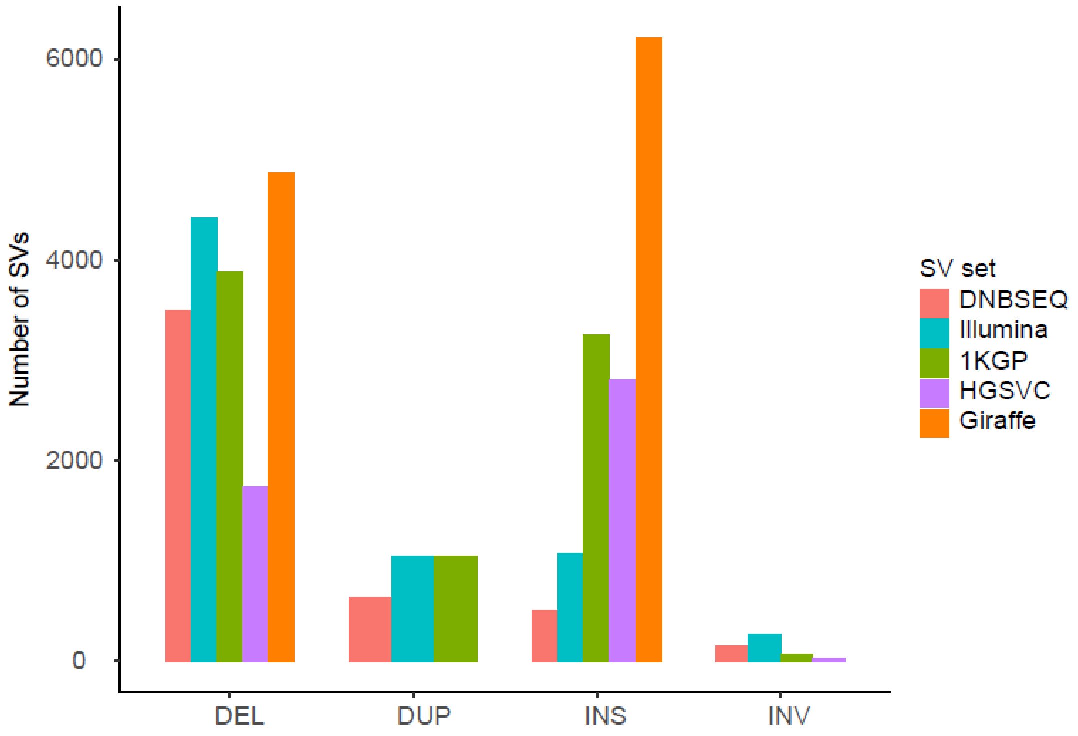


**Supplementary Figure 3. Number of SVs on different SV sets.** The bar plot presents the number of SVs across various SV types, delineated on the x-axis. The provenance of each SV set is color-coded for clarity: DNBSEQ, the DNBSEQ SV set from the current study, red; Illumina, the Illumina SV set from this study, blue; 1KGP, the SV set from Marta et al., 2022, green; HGSVC, the SV set from Peter et al., 2019, purple; Giraffe, the SV set from Jouni et al., 2021, orange.

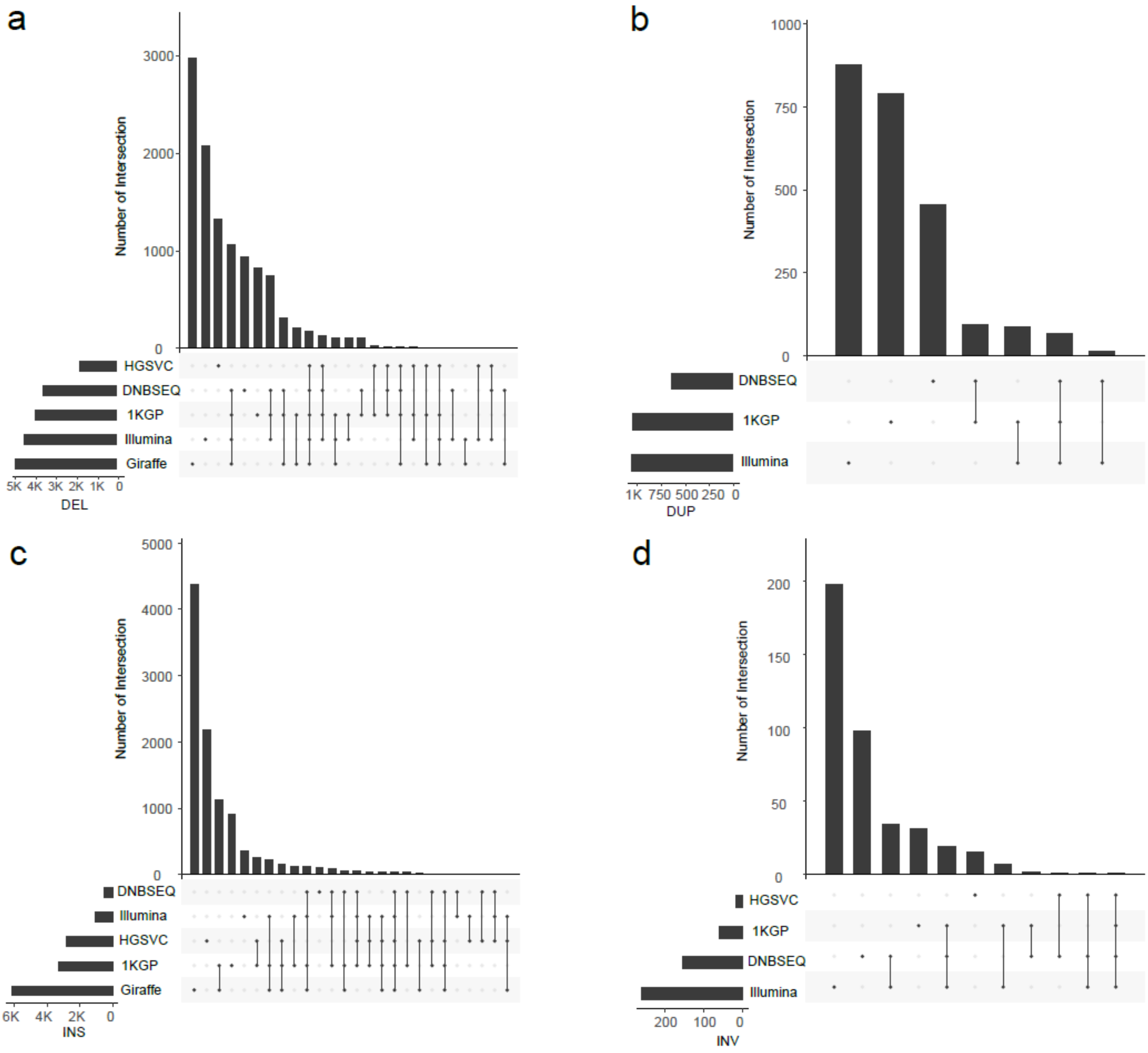


**Supplementary Figure 4. Comparison of different SV sets.** The UpSet diagram illustrates the overlap of DELs (**a**), DUPs (**b**), INSs (**c**), and INVs (**d**) across various SV sets. DNBSEQ, the DNBSEQ SV set from this study; Illumina, the Illumina SV set from this study; 1KGP, the SV set from Marta et al., 2022; HGSVC, the SV set from Peter et al., 2019; Giraffe, the SV set from Jouni et al., 2021.

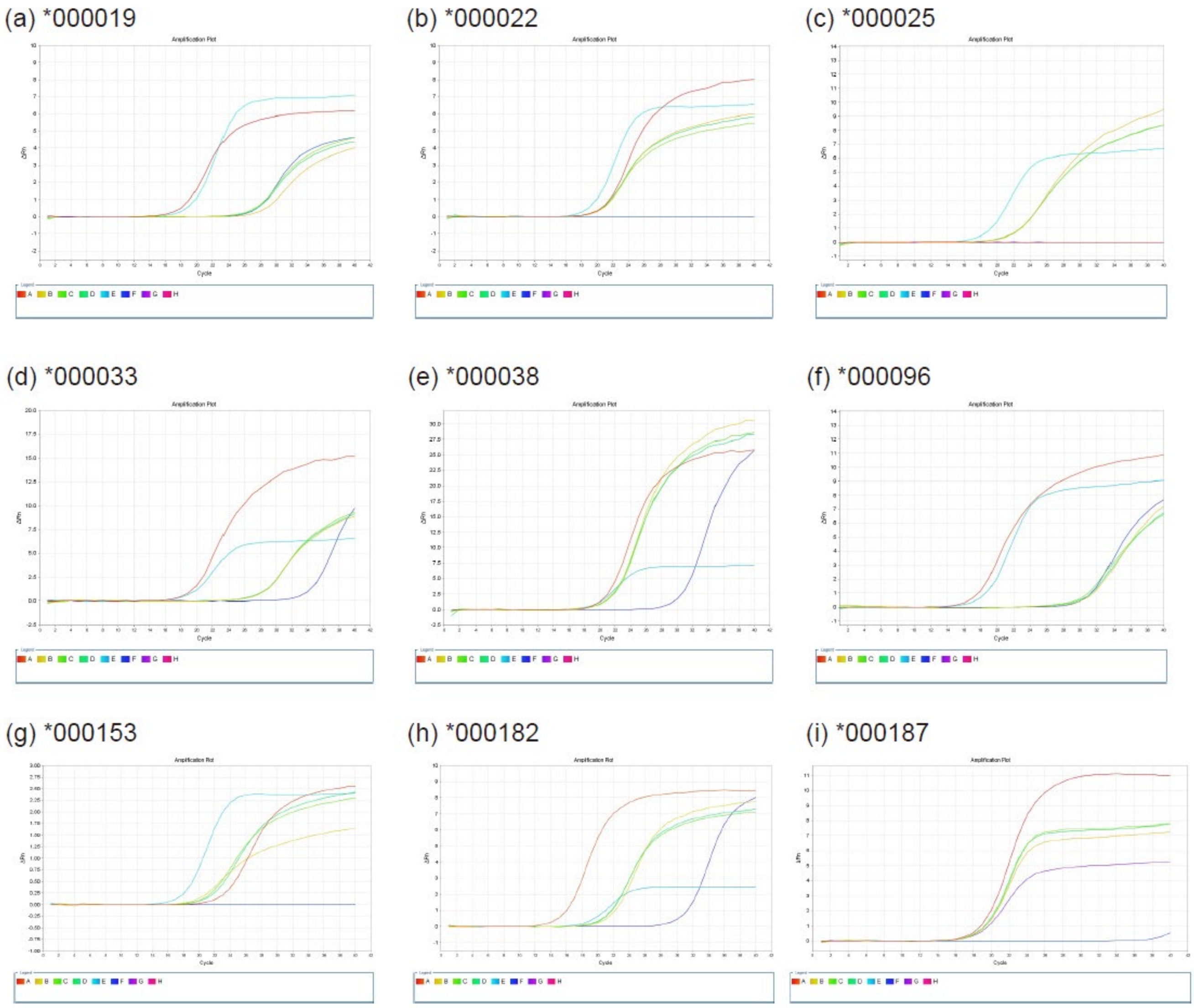


**Supplementary Figure 5. Amplification plot of DELs on real-time PCR.** Panels (**a**) – (**i**) display the results of RT-PCR experiments for nine DEL loci. In the graphs, the x-axis represents the number of cycles, while the y-axis shows the Rn value. Different colors in the graphs correspond to different samples: A: the standard sample; B-D: SV samples; E: the GAPDH housekeeping gene; and F: the negative control.

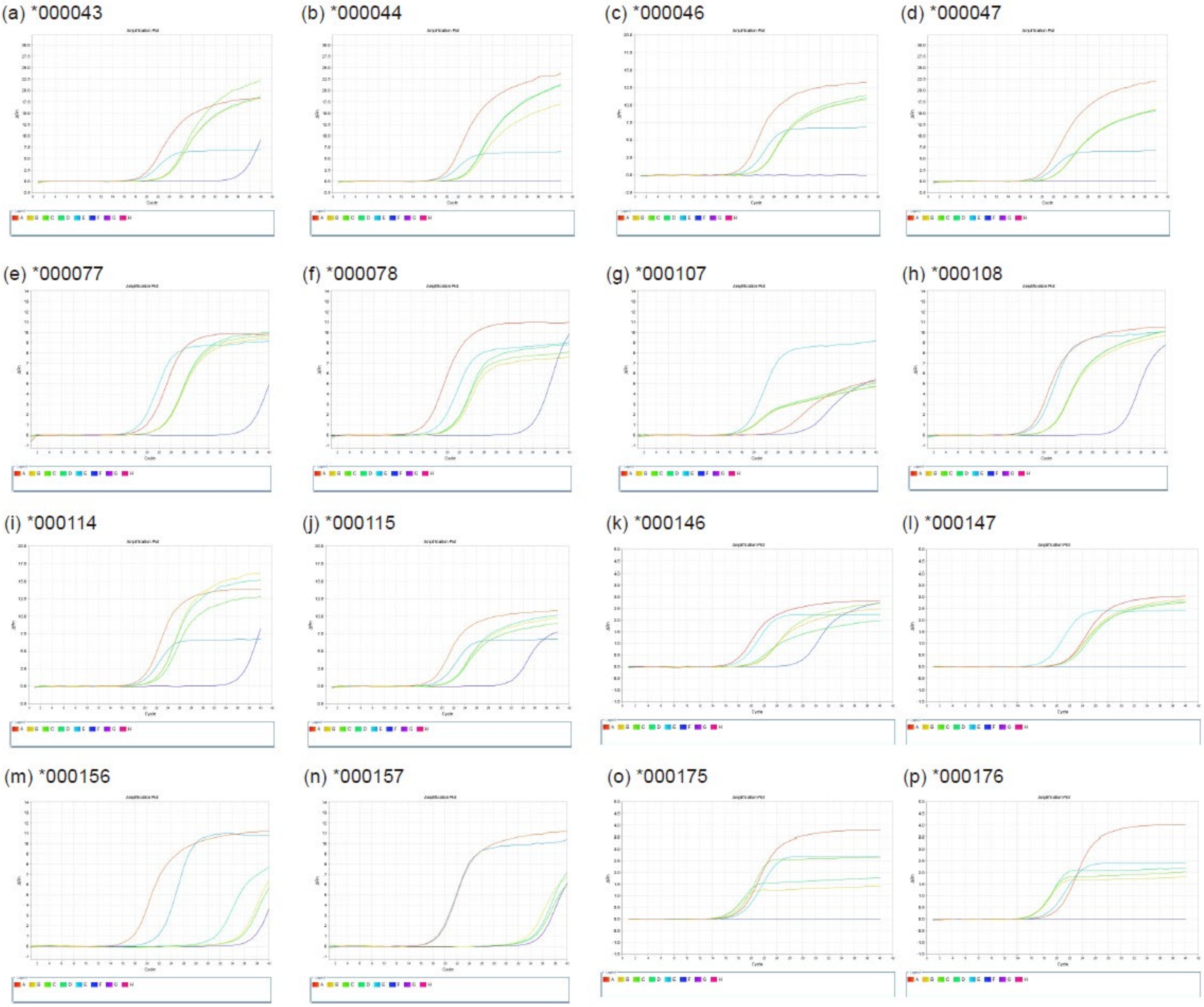


**Supplementary Figure 6. Amplification plot of INSs on real-time PCR.** Panels (**a**) – (**p**) display the results of RT-PCR experiments for 16 INS loci. In the graphs, the x-axis represents the number of cycles, while the y-axis shows the Rn value. Different colors in the graphs correspond to different samples: A: the standard sample; B-D: SV samples; E: the GAPDH housekeeping gene; and F: the negative control.

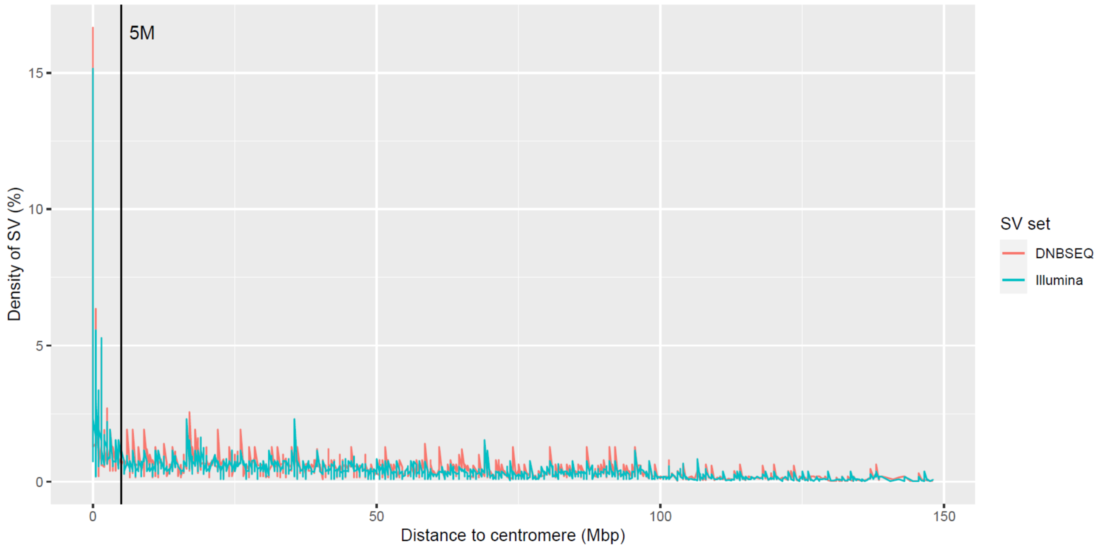


**Supplementary Figure 7. Distance to centromere.** The line graph depicts the proportion of structural variations (SVs) relative to their distance from the centromere, as indicated on the x-axis. A grey line marks the threshold at a distance of 5 Mbp from the centromere. The SV sets from DNBSEQ and Illumina are differentiated by color, with DNBSEQ in red and Illumina in blue.

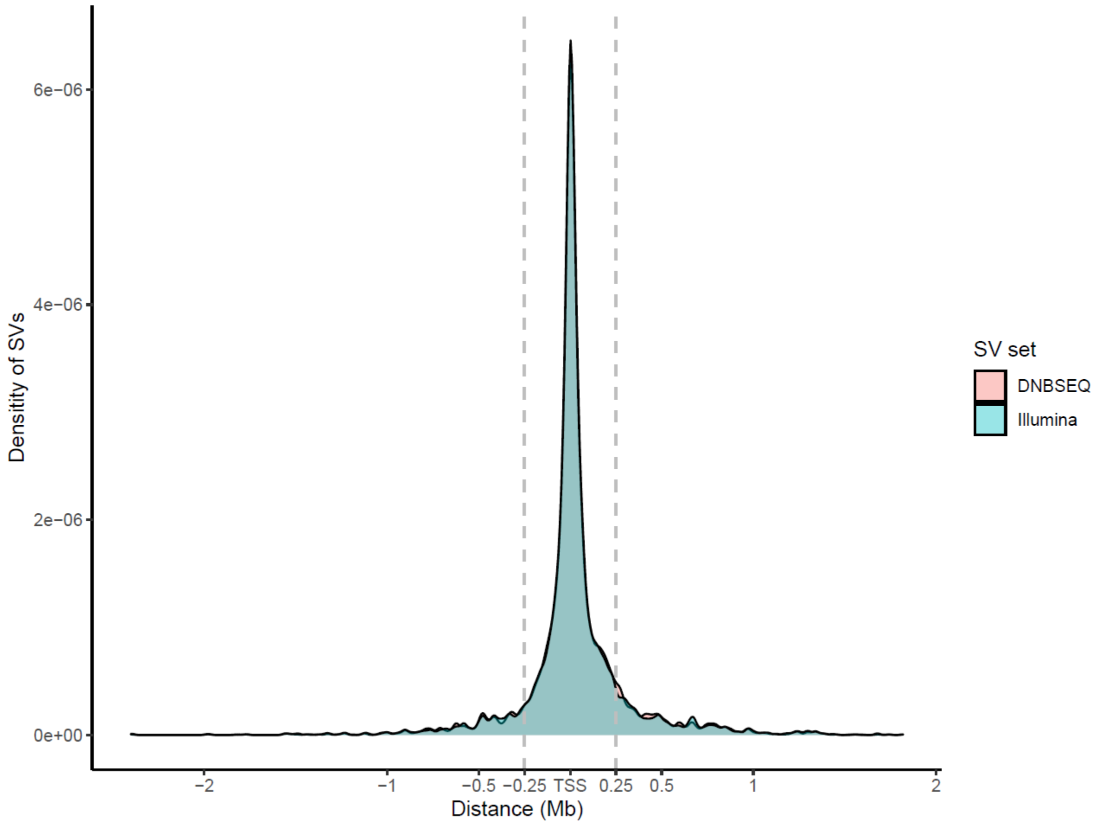


**Supplementary Figure 8. Distance to TSS.** The density plot illustrates the distribution of SVs relative to their distance from the transcription start site (TSS), as plotted on the x-axis. A grey dashed line denotes the 2.5 Mbp distance from the TSS. The SV sets from DNBSEQ are shown in red, while those from Illumina are depicted in blue.

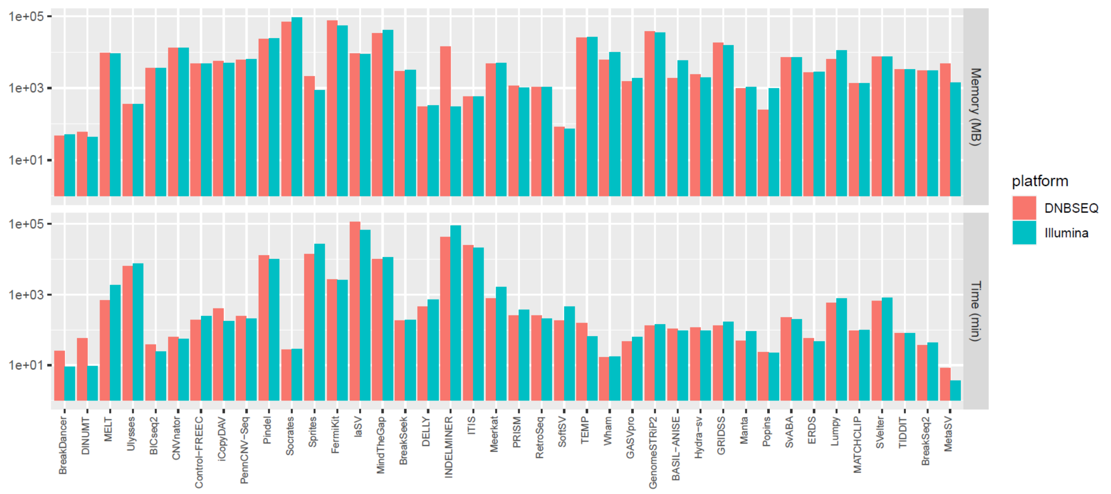


**Supplementary Figure 9. Resource Utilization in SV Detection by Different Tools.**

The bar plot quantifies the memory (**top**) and time (**bottom**) resources consumed by various tools, as indicated on the x-axis. The datasets analyzed are distinguished by color, with DNBSEQ datasets in red and Illumina datasets in blue.

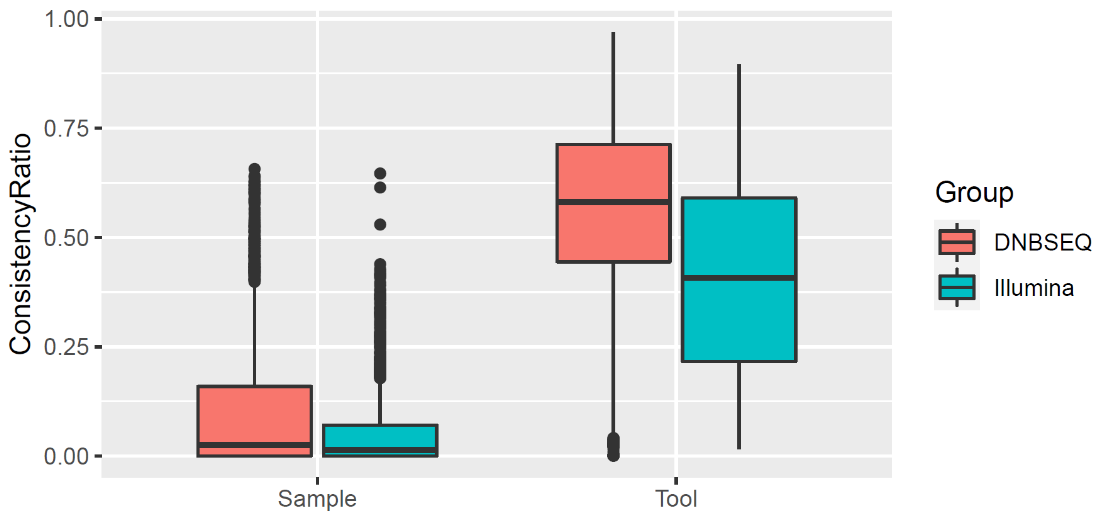


**Supplementary Figure 10. Pairwise Comparison Consistency Ratios.** The box plot depicts the consistency ratios of SV results obtained from the same dataset using different tools (labeled as 'Sample' on the x-axis) and the consistency ratios of SV results from different datasets analyzed by the same tool (labeled as 'Tool' on the x-axis). The dataset's sequencing platforms are color-coded: datasets sequenced on DNBSEQ platforms are shown in red, while those sequenced-on Illumina platforms are in blue.
